# Autotransporter folding avoids a kinetic trap during vectorial translocation across the bacterial outer membrane

**DOI:** 10.64898/2026.08.22.746429

**Authors:** Lan Yang, Qing Luan, Michael Baxa, Patricia L. Clark, James C. Gumbart

## Abstract

Autotransporter proteins are major virulence factors in Gram-negative pathogens, yet how they fold during secretion remains incompletely understood. A longstanding puzzle is why pertactin folds and is secreted in vivo within minutes but refolds in vitro over hours to days. We introduce BEAM, a multiscale framework that learns slow collective variables from coarse-grained simulations to guide all-atom enhanced sampling. Applied to a C-terminal segment of the pertactin passenger domain from *Bordetella pertussis*, BEAM achieved four- to six-fold greater conformational coverage than traditional collective-variable-guided adaptive sampling or unbiased molecular dynamics. The resulting free-energy landscape revealed a compact, non-native intermediate accessible in bulk solution but geometrically incompatible with vectorial translocation across the outer membrane. Kinetic simulations show that access to this intermediate slows folding, whereas excluding it produces rapid, in vivo-like kinetics. Together, these results explain how vectorial secretion accelerates pertactin folding by excluding an off-pathway kinetic trap. More broadly, BEAM provides a multiscale strategy for revealing hidden conformational states at atomic resolution.

## Introduction

Classical autotransporters, or Type Va secretion systems, represent the largest and most extensively studied family of secreted virulence proteins of Gram-negative bacterial pathogens, playing critical roles in pathogenesis and host-pathogen interactions^1, 2^. These proteins consist of two main functional domains: an N-terminal passenger domain that constitutes the virulence factor itself^3, 4^ and a C-terminal β-barrel domain that, with assistance from the β-barrel assembly machinery (BAM) complex, inserts into the outer membrane to facilitate passenger translocation^5–14^. The final step of autotransporter secretion involves C- to N-terminal threading of the passenger domain across the outer membrane, with folding occurring co-translocationally, a process that is both ATP and proton gradient-independent^1, 15^.

Among these systems, pertactin from *Bordetella pertussis*, the causative agent of whooping cough, serves as a paradigmatic model for understanding autotransporter biogenesis^16^. As an essential adhesin that enables bacterial attachment to respiratory epithelial cells, pertactin plays a crucial role in whooping cough pathogenesis^16, 17^. The pertactin passenger domain adopts a β-helical structure^18^ and achieves its functional conformation in the extracellular space following secretion^19, 20^. One model for ATP-independent secretion proposes that passenger folding acts as a Brownian ratchet, with extracellular folding providing directional bias^19, 21–25^. Experimental studies confirm C-to-N vectorial folding in both in vivo secretion^19^ and in vitro refolding^26^, yet kinetics differ dramatically: in vivo secretion completes within ∼20 minutes^21^ while in vitro refolding requires hours to days^20^. The molecular basis of this orders-of-magnitude kinetic discrepancy has remained a longstanding puzzle in autotransporter biology. Recently, a combination of fluorescence spectroscopy and limited proteolysis was used to show that during refolding in vitro the pertactin passenger adopts a kinetically trapped, non-native conformation with a largely native C-terminal structure and a disordered N-terminus^26^; however, the molecular basis for the energy barrier that prevents conversion of this off-pathway partially folded state (PFS) to that native state remains unknown. Understanding this difference in folding rates requires detailed, atomistic simulations that can identify conformational intermediates and folding pathways differentially accessible under vectorial versus bulk-solution conditions, ultimately providing a framework for interpreting and modulating autotransporter secretion.

Molecular dynamics (MD) simulations offer atomic-level resolution of folding processes, but studying such complex conformational transitions remains challenging even with state-of-the-art computational resources. While specialized supercomputers like Anton have enabled millisecond simulations of fast-folding proteins (typically small, α-helix-rich proteins under 100 aa)^27, 28^, other proteins like the pertactin passenger domain with β-helical structure represent a fundamentally more challenging system that requires extended simulation times far beyond the reach of conventional MD approaches. The sampling limitations encountered in studying complex protein folding are part of a broader challenge in MD simulations, which also affects the study of protein-ligand binding, allosteric conformational changes, and other processes that require crossing high energy barriers over timescales that often exceed computationally feasible ranges. To address these fundamental sampling challenges, numerous enhanced sampling methods have been developed, broadly categorized into biased and unbiased approaches^29^. Biased enhanced sampling methods apply external biasing forces to modify the underlying potential energy surface^30–32^, followed by reweighting procedures to recover the unbiased thermodynamic ensemble^33, 34^. While effective, these approaches can potentially induce unphysical conformations due to the artificial bias. In contrast, unbiased enhanced sampling methods employ strategic trajectory restart protocols based on specific criteria after initial short runs, thereby preserving the original energy landscape. Recent developments in this category include FAST^35^, AdaptiveBandit^36^, and REAP^37, 38^, among others. For a comprehensive review of these methodologies, we refer readers to the review by Kleiman et al.^39^. However, both approaches face a fundamental challenge: the selection of appropriate collective variables (CVs)—low-dimensional coordinates that capture the essential degrees of freedom governing rare transitions—that can effectively guide conformational exploration.

The identification of physically meaningful CVs becomes particularly challenging for complex systems undergoing large-scale conformational changes. Various data-driven approaches have been developed to address this challenge, with machine learning methods proving particularly effective. These include geometry-based methods (principal component analysis (PCA)^40^, diffusion map^41, 42^) that analyze con-formational similarities, committor-based approaches (transition path sampling (TPS)^43, 44^), forward flux sampling (FFS)^45^) that identify kinetic bottlenecks, and variational methods (time-lagged independent component analysis (tICA)^46, 47^, VAMPNet^48^) that identify slow dynamic modes through optimization of Markovian processes. Despite their diverse theoretical foundations, all these methods require extensive simulation data to accurately capture relevant collective motions, creating a fundamental chicken-and-egg problem: accurate CVs require extensive sampling, yet efficient sampling requires good CVs.

Current solutions to this dilemma follow two main strategies. First, some methods employ interpolation or transition-state generation techniques to construct intermediate states along putative reaction pathways^49–52^. However, these approaches often fail for complex systems with non-intuitive folding pathways. Second, adaptive-generation approaches iteratively refine CVs by running initial trajectories, generating preliminary CVs, sampling new conformations, and updating CVs until convergence^53–56^. While conceptually appealing, these methods suffer from low efficiency when initial CVs provide inadequate guidance for barrier crossing.

To address this fundamental challenge, we leverage the complementary strengths of coarse-grained (CG) and all-atom simulations. While CG models can efficiently sample large-scale conformational changes over extended timescales, they lack the atomic detail required for precise thermodynamic calculations. Here, we introduce BEAM (Boosting Enhanced sampling of All-atom simulations with Machine-learned CVs from coarse-grained trajectories), which overcomes the CV selection bottleneck by leveraging machine learning to extract effective CVs from coarse-grained simulations, enabling efficient conformational exploration at all-atom resolution. Applied to the C-terminus of the pertactin passenger domain, BEAM reveals an off-pathway intermediate that acts as a kinetic trap exclusively under in vitro conditions. This compact intermediate cannot form during vectorial secretion due to geometric constraints of the β-barrel translocon, providing a molecular explanation for the dramatic difference in folding rates between in vivo and in vitro conditions. Our findings not only advance enhanced sampling methodology but, coupled with recent, complementary experimental analyses^26^, highlight physically plausible solutions to a fundamental unresolved question in autotransporter biology, demonstrating how cellular machinery guides protein folding through spatial constraints and revealing new insights into bacterial virulence factor biogenesis.

## Results

We introduce BEAM, a workflow that uses CG simulations to generate training data for machine-learning construction of CVs, which in turn enables efficient enhanced sampling at the all-atom level. The distinguishing feature of BEAM is the transfer of slow coordinates learned from computationally efficient CG trajectories to guide adaptive sampling at all-atom resolution. A conceptual overview of BEAM and its workflow is shown in Fig. 1, with implementation details provided in the Methods. Applied to the C-terminus of the pertactin passenger domain, BEAM systematically mapped the free-energy landscape of this β-helical segment and identified key conformational intermediates along folding pathways.

**Figure 1.**
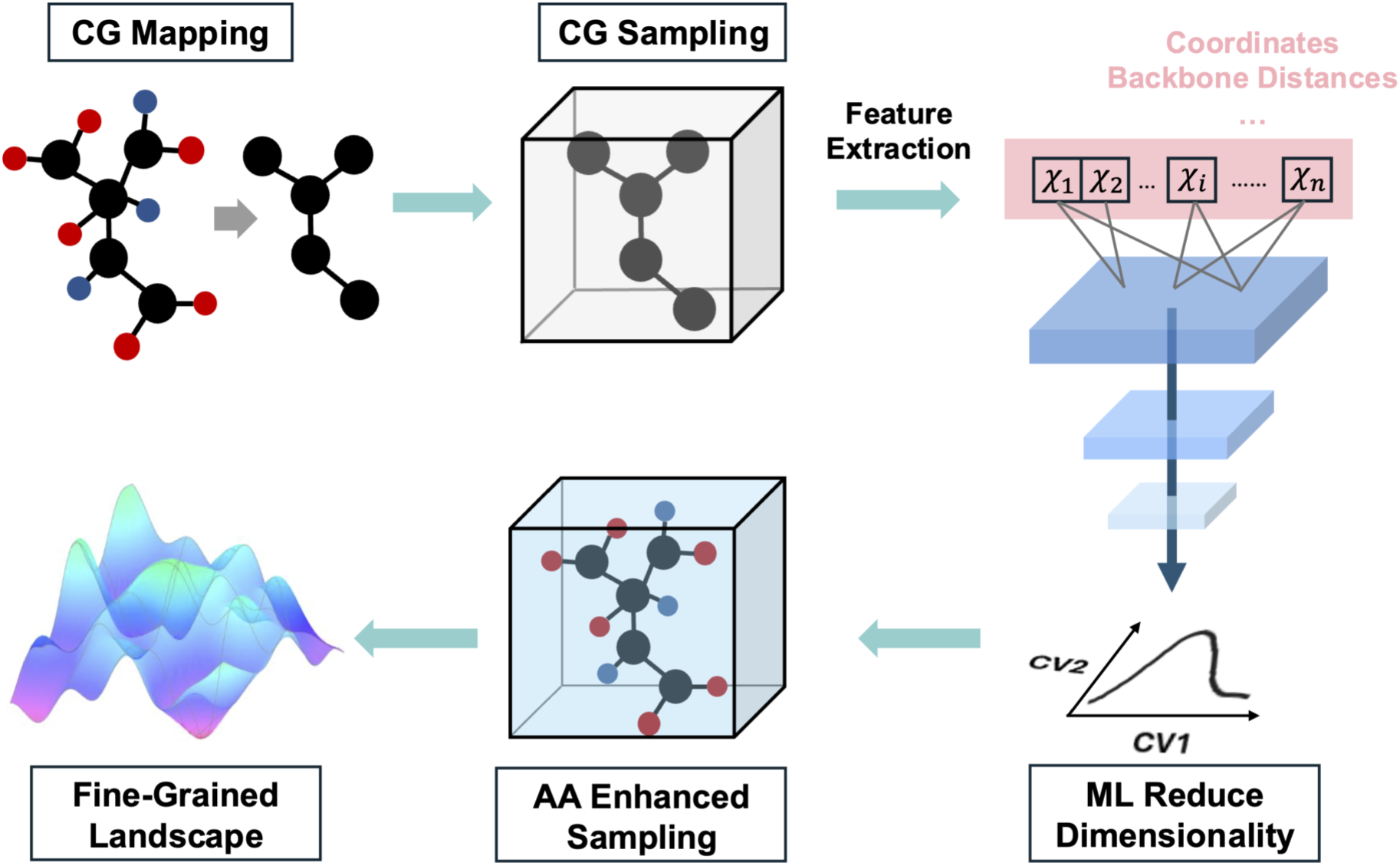
The BEAM workflow. The system is represented at CG representation to enable efficient conformational sampling. Features extracted from CG trajectories are then used in a machine-learning dimensionality-reduction step to identify CVs, which subsequently guide all-atom enhanced sampling and recover the fine-grained free-energy landscape.

### BEAM enables comprehensive sampling of pertactin unfolding landscape

We applied BEAM to the challenging task of sampling the unfolding of the C-terminal 106-residue segment of pertactin (residues 377–482), which forms an eleven-stranded β-helix (Fig. 2a). Previous steered molecular dynamics studies showed that unfolding proceeds sequentially, rung by rung, from the N terminus, due to a monotonic N-to-C stability gradient, with each successive rung becoming progressively more stable toward the C-terminus^57^. The sampling challenge of this system is not captured by residue count alone: its repeated β-helical architecture creates a high-dimensional ensemble of partially unfolded conformations. As in any folding free-energy calculation, mapping the landscape requires extensive sampling beyond native-basin fluctuations; here, that requirement is amplified by the need to cover states with different combinations of disrupted β-helical rungs, which may connect to distinct folding or misfolding routes. Such conformational heterogeneity is difficult to access with conventional MD on practical timescales, making this system a stringent and biologically relevant test case for enhanced sampling.

**Figure 2.**
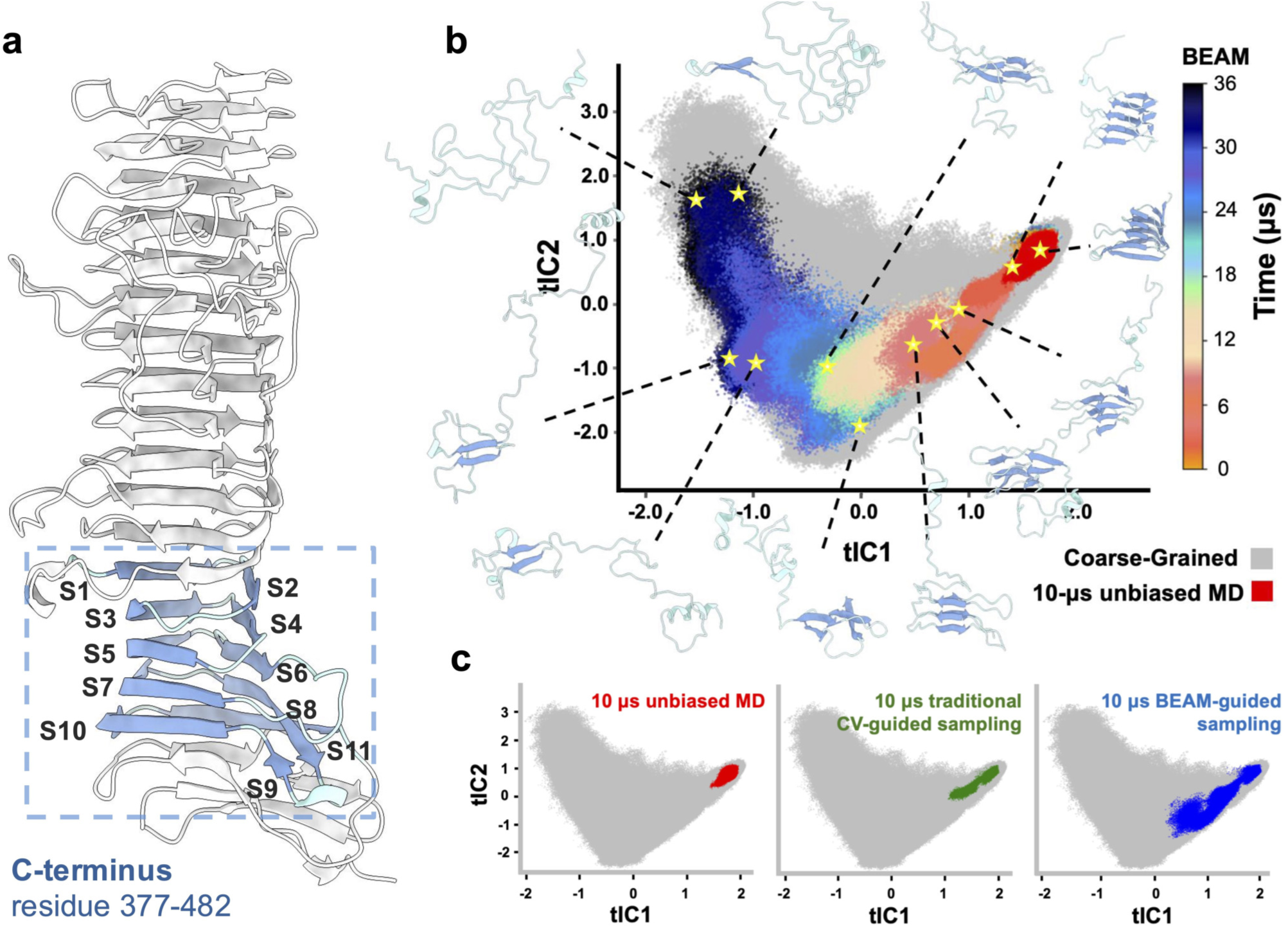
BEAM enables comprehensive conformational sampling. (a) Structure of pertactin with the simulated C-terminal segment highlighed in blue (residues 377–482, from PDB 1DAB^18^). The 11 β-strands that comprise the β-helical core studied here are labeled S1–S11 and numbered from the N-terminal side of this segment. (b) Complete conformational landscape sampled by BEAM over 36 µs, with points colored by BEAM sampling time. Selected example conformations (yellow stars) illustrate the progression from native-like folded structures to highly unfolded conformations across the sampled landscape. (c) Comparison of 10 µs sampling coverage for different methods projected onto the same tIC space: unbiased MD (left), traditional CV-guided REAP using C*_α_* –C*_α_*contacts, hydrogen bonds, radius of gyration, and RMSD (middle), and BEAM-guided REAP (right). Colored points indicate the regions sampled by each all-atom simulation, while the gray background denotes the conformational space explored by CG simulations (*Upside*)^58^.

In our implementation of BEAM, CG sampling is performed with *Upside*^59^, a machine-learned CG model that represents each residue with three backbone atoms (N, C*_α_*, C) for dynamics while inferring the coordinates of the HN, O, C*_β_*, and side chains during force calculations. *Upside* computes side-chain free energies analytically at each time step by globally computing the side-chain positional probabilities that produce the lowest free energy. This approach smooths the energy landscape by avoiding explicit side-chain dynamics, enabling efficient exploration of large-scale conformational transitions. Slow CVs are identified using time-lagged independent component analysis (tICA)^46, 47^, a dimensionality reduction method that extracts the slowest dynamical modes from trajectory data. These machine-learned CVs then guide all-atom enhanced sampling through REinforcement learning based Adaptive samPling (REAP)^37, 38^, which iteratively learns the relative importance of user-provided CVs and selects starting structures from under-sampled regions to accelerate conformational space coverage.

To evaluate the impact of these components, we compared three strategies over 10 µs: unbiased MD, REAP using traditional CVs (C*_α_* –C*_α_* contacts, hydrogen bonds, radius of gyration, RMSD), and BEAM-guided REAP using CG-derived machine-learned CVs. All trajectories were projected onto a common CV space defined by CG tICs (Fig. 2c). In contrast to the limited sampling achieved by conventional approaches, BEAM-guided REAP achieved substantially broader conformational coverage, accessing states with unfolding extending to strand S6, compared to S3 for traditional CV-guided REAP and S2 for unbiased MD. This corresponds to substantial improvements in conformational coverage: BEAM-guided REAP achieved 12.97% compared to 3.14% for traditional CV-guided REAP and 2.12% for unbiased MD, a four-fold and six-fold increase, respectively.

Given this improved sampling efficiency, we extended BEAM-guided simulations to 36 µs to obtain comprehensive coverage of the unfolding process (Fig. 2b). The resulting distribution exhibits an L-shaped topology, characteristic of systems that transition between distinct dominant modes as unfolding proceeds. Notably, the all-atom sampling (colored by simulation time) shows more limited coverage than the CG landscape (grey background), reflecting the greater energetic roughness at all-atom resolution that introduces additional barriers to conformational transitions. Nevertheless, BEAM successfully reached conformational regions that remained inaccessible to conventional approaches within comparable computational times. This extensive exploration was essential for resolving metastable intermediates and alternative unfolding pathways that help explain the stark kinetic discrepancy between in vivo secretion and in vitro refolding.

### Discovery of an off-pathway kinetic trap accessible only under in vitro conditions

To construct the free-energy landscape underlying pertactin unfolding, we performed replica exchange umbrella sampling (REUS)^60^ using all-atom–trained tIC coordinates as collective variables. These CVs were obtained by retraining tICA on BEAM-generated all-atom trajectories, ensuring consistency between the CV resolution and the simulation data used for PMF construction (see Methods). The resulting PMF reveals three dominant metastable basins (Fig. 3a): State A, the native folded state; State B, a partially unfolded intermediate occupying a distinct local basin with compact geometry; and State C, a partially unfolded intermediate with extended geometry. To quantify the geometric characteristics of these basins, we developed and computed a structural anisotropy metric for each conformation by analyzing only the unfolded residues (Fig. 3b). Anisotropy (*κ*) measures shape elongation based on the eigenvalues of each conformation’s coordinate covariance matrix (see Eq. 1 in Methods); low *κ* indicates compact, spherical shapes, while high *κ* reflects elongation. State C shows high anisotropy, consistent with extended unfolded conformations. In contrast, State B displays low anisotropy across its basin, demonstrating that it consists of compact, globular conformations rather than a mixture of geometries.

**Figure 3.**
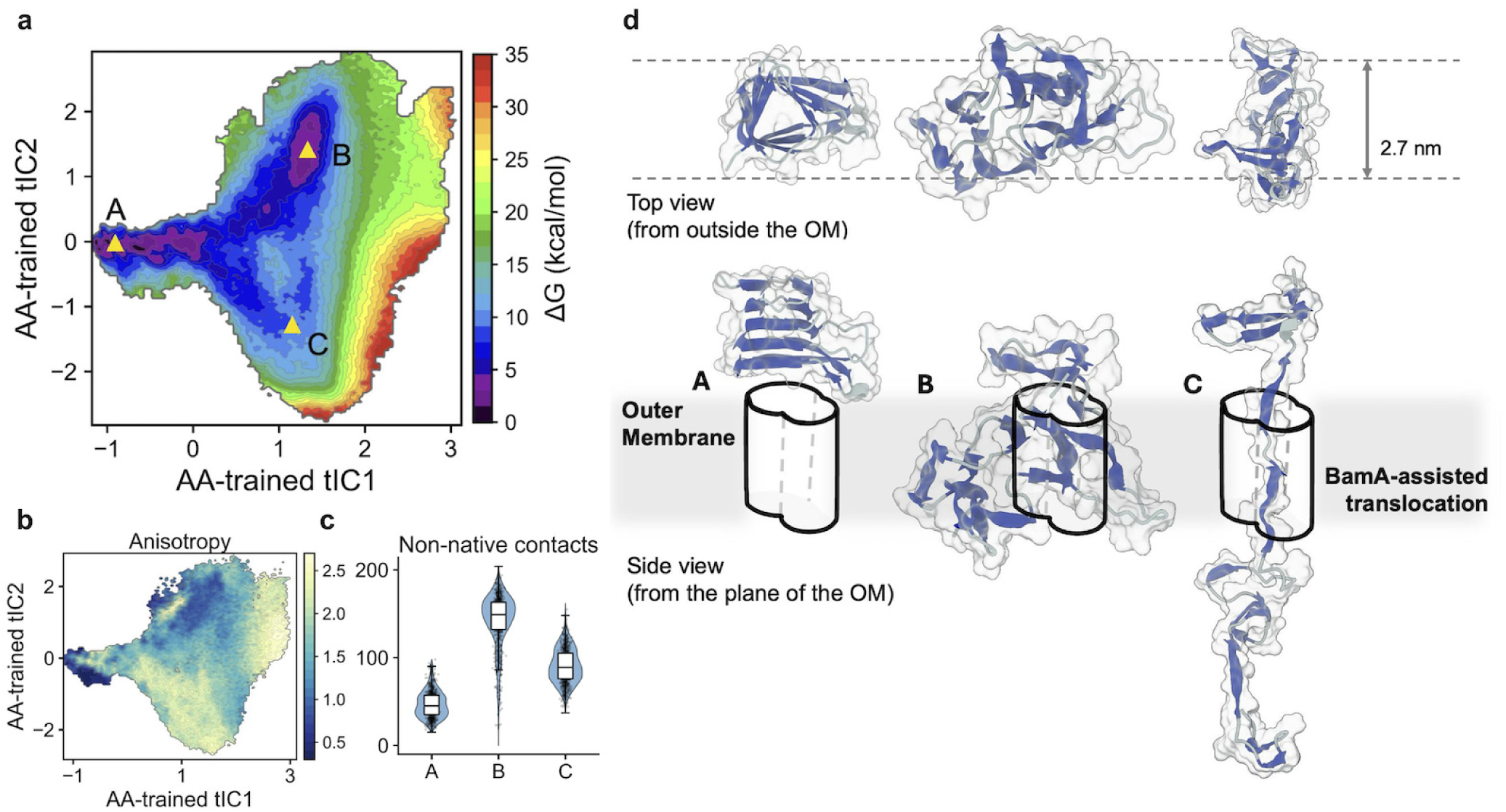
Free energy landscape reveals a compact non-native trap. (a) Potential of mean force (PMF) mapped onto all-atom-trained tIC coordinates, identifying three metastable basins: State A, the native folded state; State B, a compact intermediate; and State C, an extended intermediate. (b) Structural anisotropy (*κ*) of unfolded residues projected onto the same tIC space as in (a). Low *κ* indicates compact geometries, whereas high *κ* indicates elongated geometries. (c) Distribution of non-native intramolecular contacts within the simulated passenger segment for frames assigned to States A–C. Violin plots show frame-wise distributions, with box plots indicating the median and interquartile range. State B is enriched in non-native contacts, consistent with a compact, non-native collapsed ensemble. (d) Schematic representation of States A–C in the context of the BamA-assisted translocon. Top views from outside the outer membrane are shown above, with side views from the plane of the outer membrane below. Dashed lines indicate the 2.7 nm reference width used for the schematic translocon. State B is incompatible with sequential vectorial translocation and sequential folding, whereas State C remains compatible with vectorial threading.

Importantly, the low free energy of State B should not be interpreted as native-like stability of a single representative structure. The PMF reflects the population of an ensemble projected onto the two tIC coordinates. To further characterize this ensemble, we quantified non-native intramolecular contacts for frames assigned to States A–C (Fig. 3c). State B is strongly enriched in non-native contacts compared with States A and C, supporting its assignment as a compact, non-native collapsed ensemble rather than preservation of the native β-helical fold. Such compaction is disfavored during vectorial translocation, where short segments emerge and fold sequentially onto the growing β-helix.

Structural analysis reveals that this compact geometry renders State B incompatible with passage through the membrane translocon (Fig. 3d). During autotransporter secretion, the passenger domain is threaded through a channel formed by the autotransporter β-barrel in conjunction with BamA^9, 61, 62^. Although the BamA-assisted autotransporter hybrid pore has not been assigned a single well-defined diameter, previous studies provide approximate geometric constraints. Mature autotransporter β-domain pores have an inner diameter of ∼1.0 nm, whereas the active autotransporter translocation pore has been functionally estimated to reach up to ∼1.7 nm^63^. Open BamA structures show an anisotropic exit pore of ∼1.5 × 2.7 nm^13^, and BamA–EspP structures support an expanded, triangular hybrid-barrel intermediate rather than a circular pore^61^. For visualization, we therefore used the long-axis dimension of the open BamA exit pore (∼2.7 nm) as a generous upper-bound scale for the schematic BamA-assisted translocon, while treating the pore as an anisotropic hybrid opening rather than an additive combination of two complete barrels. State B consists of compact conformations with uniformly low anisotropy, reflecting near-spherical geometries that are incompatible with threading through the confined channel. In contrast, the elongated conformations of State C, characterized by high anisotropy, can readily thread through during vectorial translocation across the outer membrane. These geometric constraints establish State B as an off-pathway kinetic trap accessible only under in vitro folding conditions, where the protein folds in bulk solution without spatial confinement.

### Kinetic simulations validate the role of State B as a folding trap

To directly test the hypothesis that State B acts as a kinetic trap, we performed kinetic Monte Carlo simulations using the Gillespie algorithm on the REUS-derived free-energy landscape (see Methods). We simulated 1000 independent folding trajectories starting from fully unfolded, extended conformations and calculated the percentage of successful folding events as a function of time under two conditions: with State B accessible (trap involved) or with State B blocked (trap blocked), as shown in Fig. 4a. In the trap-blocked condition, grid cells corresponding to the State B basin were removed from the accessible kinetic network.

**Figure 4.**
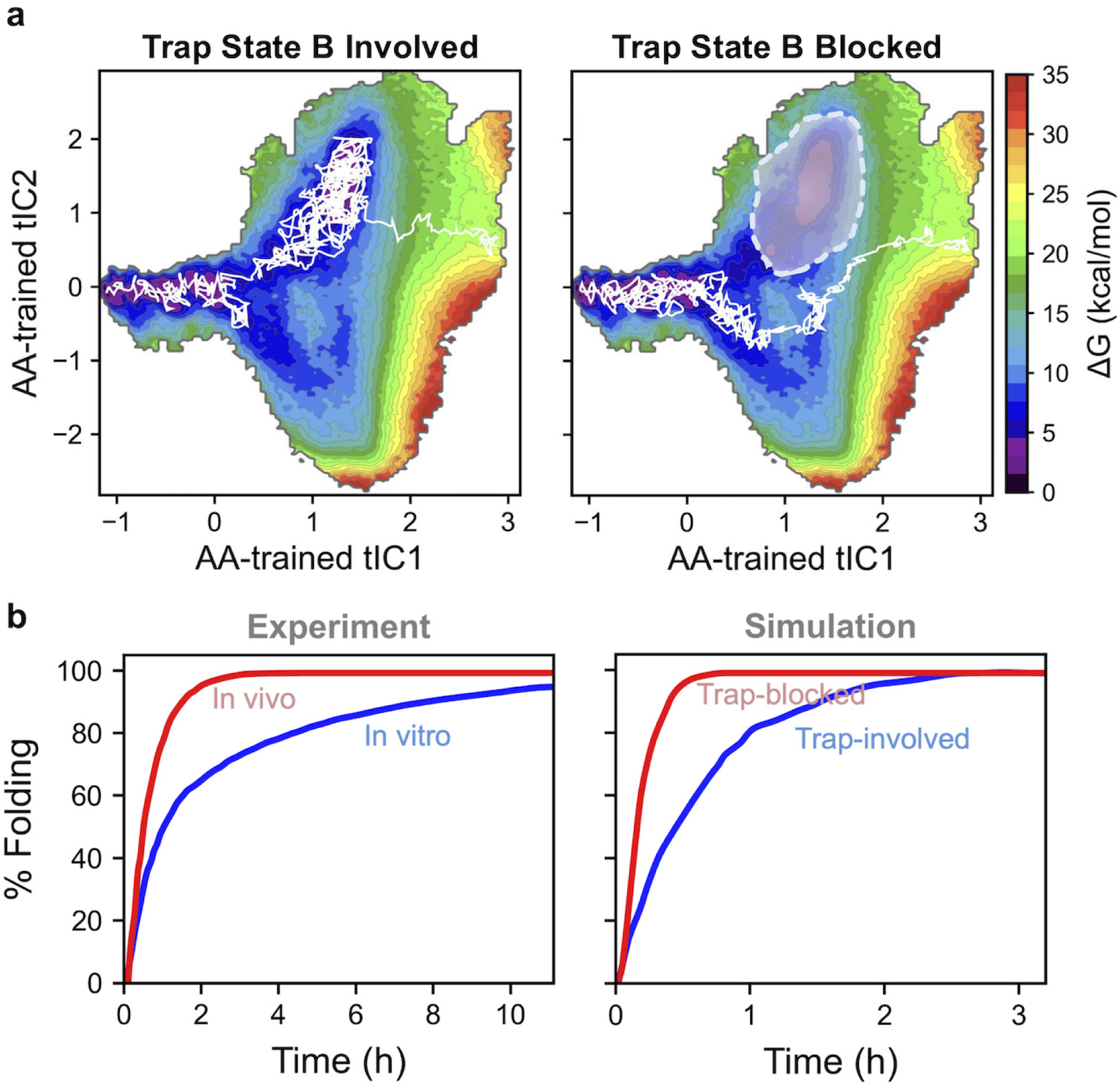
Kinetic simulations validate State B as a folding trap. (a) Schematic representation of folding pathways with State B accessible (left) or blocked (right). The PMF landscape shows the three metastable states, with the blocked region indicated in the shaded area in right panel. White trajectories illustrate representative folding routes from unfolded regions to the native state A. (b) Left: Experimental folding kinetics of pertactin passenger domain, comparing in vivo secretion (red) and in vitro refolding (blue), adapted from Ref.^21^. Right: Gillespie algorithm simulations of 1000 independent folding trajectories with State B accessible (blue) or blocked (red), demonstrating accelerated folding when the kinetic trap is eliminated.

Our kinetic simulations with State B accessible (Fig. 4b, blue curve) qualitatively reproduce the slow timescale observed experimentally for in vitro refolding, while blocking State B (Fig. 4b, red curve) leads to substantially faster folding. Several factors limit quantitative comparison with experiment. First, our simulations model only the C-terminal 106-residue segment, whereas experimental measurements reflect folding of the complete ∼ 500-residue passenger domain. The additional N-terminal segments likely introduce further kinetic complexity not captured in our model. Indeed, a recent study of the folding of the C-terminal ∼200 aa of the pertactin passenger found that both the presence and the foldability of N-terminal segment (∼300 aa) created kinetic traps for the folding of the C-terminus, yet the folding rate of the ∼200 aa C-terminal segment alone was still significantly slower than the estimated slowest possible time scale of protein folding in vivo^26^, suggesting additional energy barrier(s) also exists within the passenger C-terminus. Second, the Gillespie algorithm employs an effective diffusion coefficient derived from mean-square displacement analysis, which provides only approximate timescale estimates. Third, detailed kinetic curve shapes depend sensitively on the distribution of starting conformations, which may differ between our extended initial states and the experimentally denatured ensemble.

Despite these limitations, our key conclusion, that State B acts as a kinetic trap, relies on relative rather than absolute folding timescales. Folding is markedly accelerated when State B is blocked, whereas allowing access to State B slows folding and produces kinetics qualitatively consistent with slow in vitro refolding. Additional intermediates present in the full 500-residue passenger domain likely contribute further to the remaining kinetic discrepancy. For example, an on-pathway intermediate was identified for the ∼500-residue full length pertactin passenger that exists at the junction between folding to the pertactin native structure versus formation of an off-pathway folding intermediate that is kinetically trapped^26^. Given the modular, repeating structure of the pertactin passenger β-helix, it is possible that similar kinetic competitions exist for other portions of the β-helix, including the C-terminal 106-residue segment studied here, and shape the overall energy landscape for folding. The geometric incompatibility of State B with vectorial secretion provides a further explanation for why in vivo folding is dramatically faster than refolding in bulk solution.

### Physical interpretation of collective variables reveals sequential unfolding mechanism

To interpret the all-atom-trained tICs used in the PMF and kinetic analyses, we analyzed their tICA weight vectors across the protein structure. Because each tIC is a linear combination of backbone atom Cartesian coordinates, the three coefficients associated with each atom define a 3D weight vector. Displacement of an atom along this vector increases the corresponding tIC value, whereas the vector magnitude reports the relative contribution of that atom to the mode. At the residue level, we computed the average weight-vector magnitude over the three backbone atoms (N, C*_α_*, C) of each residue for tIC1 and tIC2 (Fig. 5b). Both tICs show oscillating loading patterns that align with native *β*-strand positions, with larger contributions in strand regions and smaller contributions in connecting loops. Superimposed on this strand-level pattern, the loadings increase toward the C-terminal rungs, consistent with the *β*-helix stability gradient: N-terminal rungs unfold earlier, whereas the more stable C-terminal rungs dominate the slow collective modes associated with late-stage unfolding.

**Figure 5.**
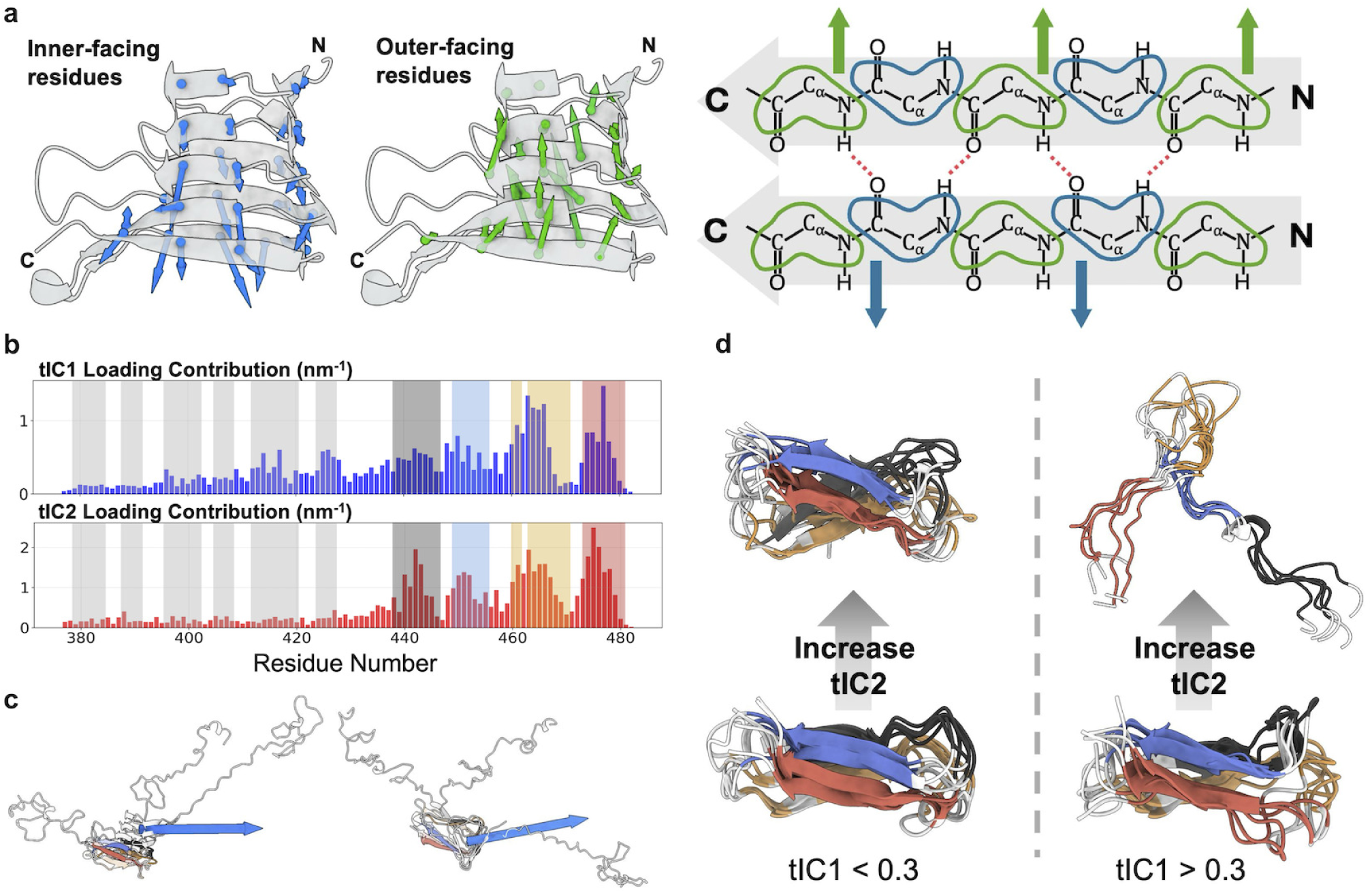
Physical interpretation of collective variables. (a) Structural interpretation of tIC1. Left: residue-averaged tIC1 weight vectors mapped onto the β-helix, showing inner-facing residues (blue) and outer-facing residues (green) exhibit consistent opposite directional patterns. Right: schematic of the right-handed β-helix geometry explaining why inward-facing and outward-facing residues must move in opposite directions during unfolding to disrupt inter-rung hydrogen bonds. (b) Residue-wise loading magnitudes of tIC1 (blue, top) and tIC2 (red, bottom). Gray shaded regions indicate native β-strands S1–S6 along the sequence. Colored shaded regions mark four terminal structural units within β-strands S7–S11 that share loading peaks; the short S9 strand is grouped with S10 because their loading peaks overlap. The same structural-unit color scheme is used in panels (c) and (d). (c) Average tIC2 motion over residues 377–436, shown in top and side views. Blue arrows indicate increasing tIC2. (d) Representative structures from two tIC1 regimes. Within each regime, tIC2 increases from bottom to top. Across the transition around tIC1 = 0.3, tIC2 shifts from capturing local rearrangements of largely folded terminal rungs to capturing terminal-rung unfolding and expanded conformational heterogeneity.

tIC1 tracks the degree of *β*-helix unfolding. To quantify this relationship, we measured the average C*_α_* –C*_α_* distances between corresponding stacked residues in neighboring *β*-helix rungs and projected these distances onto the (tIC1, tIC2) space (Figure S1). These distances increase systematically with tIC1, indicating that larger tIC1 values correspond to separation of neighboring rungs and opening of the *β*-helical structure. Therefore, the direction of each tIC1 weight vector can be interpreted as the direction in which that residue moves as the structure progresses toward more unfolded conformations. Mapping these residue-averaged tIC1 weight vectors onto the protein structure reveals a face-dependent directional pattern (Fig. 5a): inner-facing and outer-facing residues exhibit opposite vector components along the long axis of the *β*-helix, oriented from the N- to C-terminal end of the simulated segment. This opposing axial motion reflects the geometric constraints of the right-handed *β*-helix and provides a structural mechanism for separating neighboring rungs and disrupting inter-rung hydrogen bonds during unfolding.

tIC2, in contrast to tIC1, represents a faster and more localized mode. Its loading pattern is bimodal: residues 377–436 carry smaller but coordinated weights, whereas the terminal region contains four dominant loading peaks within strands S7–S11 (Fig. 5b). Averaging the tIC2 vectors over residues 377–436 reveals a consistent directional bias toward increasing tIC2 (Fig. 5c), while the larger terminal loadings prompted us to examine the structural behavior of the C-terminal rungs directly.

To summarize this behavior, we selected representative structures from two tIC1 regimes separated at tIC1 = 0.3 and compared how the terminal region changes as tIC2 increases (Fig. 5d). Below this boundary, the terminal rungs remain largely folded and tIC2 mainly captures local rearrangements. Above this boundary, terminal-rung unfolding expands the accessible tIC2 range, and tIC2 increasingly reflects broader conformational diversity. Thus, tIC2 is a mixed-mode descriptor: it combines a directional bias from residues 377–436 with large-amplitude rearrangements of the terminal rungs that become prominent during late-stage unfolding. The CG-derived tICs show qualitatively consistent patterns with these all-atom-trained tICs, supporting the multiscale sampling strategy.

## Discussion

In this work, we introduced BEAM, a multiscale computational framework that addresses the central challenge of collective variable (CV) generation for enhanced sampling by leveraging CG simulations to guide all-atom exploration. By training machine learning models on computationally accessible CG trajectories, BEAM circumvents the data scarcity that traditionally limits data-driven CV discovery in biomolecular systems. The resulting multiscale workflow combines the sampling efficiency of CG models with the structural accuracy of all-atom simulations, enabling the exploration of conformational landscapes that remain inaccessible to conventional approaches within comparable computational budgets.

Applied to the pertactin passenger domain, BEAM revealed a compact, collapsed intermediate (State B; Fig. 3a) that forms exclusively under in vitro conditions and acts as a kinetic trap. The compact geometry of this intermediate is inaccessible during vectorial translocation and sequential folding across the outer membrane in vivo, leading to significantly faster folding than bulk refolding. By constraining accessible conformational space, the secretion machinery effectively eliminates off-pathway intermediates, offering a molecular explanation for the long-standing kinetic discrepancy in autotransporter folding. This mechanism may extend to other autotransporter virulence factors and to broader folding processes influenced by geometric confinement, including cotranslational folding within the ribosome exit tunnel and, more generally, chaperonin-assisted folding in GroEL/GroES.

BEAM’s reliance on CG simulations assumes that the essential transitions and physical interactions are preserved at reduced resolution. This assumption is well satisfied for large-scale backbone-dominated motions such as β-helix unfolding but may break down for processes driven by side-chain specificity, electrostatics, or local frustration, e.g., allosteric transitions or binding-pocket rearrangements. Furthermore, selecting an appropriate CG model remains largely empirical, requiring case-by-case assessment of whether the chosen representation adequately captures both the relevant kinetics and the sampling efficiency. Extending BEAM to additional autotransporters, as well as to proteins undergoing large-scale folding, domain motions, or aggregation, will require careful selection of an appropriate CG model and validation that the reduced representation faithfully captures the key physical driving forces that are relevant to the conformational transitions of interest.

Despite these limitations, BEAM offers a practical and general strategy for overcoming a fundamental bottleneck in enhanced sampling: identifying informative collective variables when intuitive reaction coordinates are unavailable. By using a computationally accessible level of resolution to bootstrap sampling at atomic detail, BEAM enables the discovery of hidden intermediates and mechanistic pathways that are difficult to detect experimentally or computationally. Our results highlight how multiscale modeling can bridge in vitro and in vivo protein folding behaviors and provide a framework for studying conformational processes shaped by geometric, cellular, or vectorial constraints.

## Methods

### BEAM workflow implementation

In this study, BEAM was implemented by first performing CG simulations to efficiently explore large-scale conformational transitions. CG trajectories were then featurized and subjected to time-lagged independent component analysis (tICA) to identify slow collective variables, which were subsequently used to guide all-atom enhanced sampling.

CG simulations were performed using *Upside* with 300 unfolding replicas, yielding 252 µs of CG sampling spanning conformations from the native fold to fully extended states. CG trajectory frames were aligned and featurized using backbone atom coordinates. tICA was performed using PyEMMA^64^ with a 4 time of 4 ns (100 Upside time steps; 1 Upside time unit = 40 ps). The choice of lag time was validated by performing a lag-time scan over 0.4–200 ns and projecting the CG conformational ensemble onto the first two tICs at each lag time. All lag times from 2-40 ns showed consistent separation of folded and unfolded states with continuous transition paths (Fig. S2). A lag time of 4 ns was selected as a compromise between statistical robustness at shorter lag times and adequate separation of slow conformational modes.

All-atom enhanced sampling was conducted using REAP, with CG-derived tICs supplied as CVs. The first two tICs, corresponding to the slowest dynamical modes, were selected as collective variables for guiding all-atom sampling for several reasons. First, tICA orders collective variables by decreasing dynamical timescale, so the leading tICs capture the slowest, most mechanistically relevant motions for large-scale conformational transitions. Second, although additional tICs capture progressively faster modes, including them would expand the CV space and distribute sampling effort across less relevant dimensions, reducing the efficiency. Specifically, REAP’s adaptive sampling protocol requires clustering conformations in CV space and dynamically reweighting CVs to prioritize under-sampled regions; both operations become significantly more expensive in higher dimensions without commensurate gains in sampling efficiency when the dominant slow modes are already captured. Finally, the two-dimensional representation also facilitates direct visualization of the conformational landscape and interpretation of folding pathways. An initial 10-µs unbiased MD trajectory seeded the conformational pool. Iterative REAP rounds launched ten 10-ns replicas per round, with starting structures selected to prioritize under-sampled regions of the CV space. Across all rounds, 36 µs of all-atom data were accumulated.

### Free-energy landscape construction

We employed REUS to obtain an all-atom PMF for the pertactin folding landscape. The BEAM-guided enhanced sampling progressed unidirectionally from folded to unfolded states without reaching global equilibrium, making direct PMF reconstruction from these trajectories impossible. Although one-directional enhanced sampling can yield kinetic information through frameworks such as history-augmented Markov state models (haMSMs)^65^, which require only local equilibrium, such approaches provide rates only in the sampled direction. Because BEAM primarily captured unfolding trajectories while our biological question concerns folding kinetics underlying the in vivo/in vitro discrepancy, haMSM-based reconstruction is insufficient. REUS instead provides the equilibrium sampling required for accurate PMF estimation.

To construct the REUS collective variables, we retrained tICA coordinates using the all-atom unfolding trajectories generated by BEAM. As in the CG stage, the aligned Cartesian coordinates of backbone atoms served as input features; a lag time of 10 ns was used to extract the slowest dynamical modes, yielding two tIC coordinates at all-atom resolution. The 10-ns lag time was determined by the length of individual REAP trajectory segments. This retraining ensures that the CVs faithfully capture relevant conformational motions at atomic detail, and these all-atom tICs were used as the reaction coordinates for REUS. BEAM’s extensive conformational sampling provided optimal initial structures for the REUS windows—physically realistic configurations drawn directly from equilibrium MD rather than artificially generated structures from interpolation or steered MD. The two-dimensional CV space was discretized into a regular grid with spacing of 0.08 tIC units along both tIC1 and tIC2, resulting in 1776 umbrella windows. Each window employed a harmonic restraint with a force constant of 50 kcal/mol/(tIC unit)^2^ to maintain sampling within the designated region. All windows were simulated for 2.2 ns, producing a total of ∼3.9 µs of REUS sampling. PMF convergence was confirmed by monitoring the RMSD between cumulative and final free-energy surfaces; RMSD stabilized below 0.5 kcal/mol after 1.5 ns per window (Fig. S3). Window placement was guided by the broad conformational coverage achieved by BEAM, ensuring that all relevant regions of the folding landscape were represented. The PMF was constructed using the weighted histogram analysis method (WHAM)^66^, which combines statistics from all windows into a continuous free-energy surface defined over the two collective variables.

### Anisotropy calculation

To quantify the geometric properties of unfolded conformations across the free-energy landscape, we computed a structural anisotropy metric, *κ*, for each trajectory frame.

Unfolded residues were identified on a per-frame basis by monitoring native C*_α_* –C*_α_*contact pairs (Table S1); a contact was considered broken when the corresponding distance exceeded 0.8 nm, and residues participating in broken contacts were classified as unfolded.

For frames containing at least three unfolded residues, the Cartesian coordinates of the corresponding C*_α_* atoms were extracted, centered by removing the centroid, and used to construct a coordinate covariance matrix. Diagonalization of this matrix yielded three eigenvalues (*λ*_1_ ≥ *λ*_2_ ≥ *λ*_3_), from which anisotropy was defined as

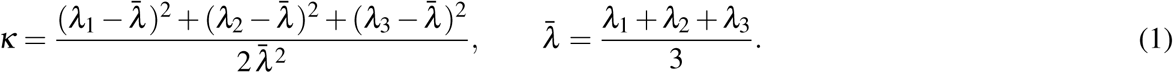

Low anisotropy corresponds to compact, near-spherical conformations, whereas high anisotropy reflects elongated geometries.

To visualize anisotropy across the conformational landscape, *κ* values were projected onto the (tIC1, tIC2) space using a regular grid with 0.05-unit spacing. For each grid cell, anisotropy was averaged over all frames within the cell, excluding cells containing fewer than five frames to suppress statistical noise from sparsely sampled regions and reduce the influence of outlier conformations. The resulting grid-averaged anisotropy map is shown in Fig. 3b.

### Kinetic simulations

To quantify folding pathways on the REUS-derived energy landscape, the PMF was discretized onto the same 0.08-unit grid. Only grid points with converged free-energy estimates were retained as microstates and connected to their eight nearest neighbors.

Position-dependent diffusion coefficients *D*(tIC1, tIC2) were estimated using the mean-square displacement (MSD) approach of Hummer^67^. We computed the local MSD for each grid point by pooling all unbiased MD trajectory segments generated during REAP, defined as MSD(*τ*) = ⟨|tIC(*t* + *τ*) −tIC(*t*)|^2^⟩ with *τ* = 2 ns. For each grid cell, all trajectory frames falling within that cell were collected, and the MSD was computed between frames separated by 2 ns. The local diffusion coefficient was then extracted using *D* = MSD(*τ*)/(2*dτ*), where *d* = 2 is the dimensionality of the tIC space. Grid cells with insufficient sampling (< 20 frames) were assigned diffusion coefficients interpolated from neighboring cells.

Transition rates were computed using a discretized Smoluchowski formulation^68^,

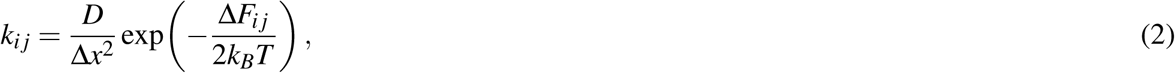

where Δ*x* = 0.08 is the grid spacing, Δ*F_i j_* is the free-energy difference, and *D_ij_*= (*D_i_* + *D_j_*)/2 is the arithmetic mean of the local diffusion coefficients. Folding trajectories were simulated using the Gillespie algorithm^69^, with 1,000 independent runs initiated from extended states. To assess the impact of specific intermediates, simulations were repeated with selected regions removed from the accessible state space to mimic geometric exclusion during vectorial secretion.

### Molecular dynamics simulation details

The pertactin C-terminal domain (residues 377–482; PDB: 1DAB) was solvated in a 16.0 × 7.0 × 7.0 nm^3^ TIP3P water box^70^ containing 0.15 M NaCl (∼ 74,000 atoms). Simulations were performed in NAMD 2.13^71^ using the CHARMM22* force field^72^, with temperature maintained at 310 K (Langevin dynamics) and pressure at 1 atm (Langevin piston). Long-range electrostatics were computed with PME (1.2-nm cutoff), and van der Waals interactions used a switching function from 1.0 to 1.2 nm. A 2-fs timestep was used with constrained hydrogen bonds.

A harmonic orientational restraint was applied using the orientation collective variable in the NAMD Colvars module^73^. The restraint was applied to the C*_α_*atoms of residues 377–482 to maintain the orientation of the protein relative to a reference structure aligned with the long dimension of the simulation box, thereby accommodating extended unfolded conformations.

For REAP adaptive sampling, the conformational pool was clustered in CV space using k-means clustering with *k* = 2,000. Within the REAP selection workflow, the top 200 cluster representatives were retained as candidate starting structures, and 10 structures were selected to launch replicas in each sampling round.

## Data Availability

Simulation input files, structures, and source data for all figures are available on Zenodo at https://doi.org/10.5281/zenodo.22035221. The experimentally determined pertactin structure used in this study is available from the Protein Data Bank under accession code 1DAB.

## Code Availability

The source code implementing BEAM, together with scripts used for the analyses presented in this study, is available on GitHub at https://github.com/LanYang430/pertactin-beam, and has been permanently archived on Zenodo at https://doi.org/10.5281/zenodo.22019935.

## Author Contributions

L.Y. performed the computational work and analyzed the data. Q.L., M.B., and P.L.C. contributed to interpretation of the results. J.C.G. conceived and supervised the study. L.Y. and J.C.G. wrote the manuscript with input from all authors. All authors reviewed and approved the final manuscript.

## Acknowledgements

This work was supported by the National Institutes of Health (NIH; R01-GM148586) and in part by the CSSE@GT Ph.D. Fellowship to L.Y. Computational resources included the Phoenix cluster, which is managed by the Partnership for an Advanced Computing Environment (PACE) at the Georgia Institute of Technology.

