## Supporting Information for "Autotransporter folding avoids a kinetic trap during vectorial translocation across the bacterial outer membrane"

### 1 Native Cα–Cα Contact Distances Track the Global Unfolding Coordinate

Native C$\alpha$–C$\alpha$ contact pairs were defined from the folded $\beta$-helix structure based on inter-rung hydrogen-bond ladders. For each trajectory frame we computed the mean distance over all native contact pairs. When projected onto the tIC1–tIC2 landscape, this mean distance increases monotonically with tIC1, confirming that tIC1 primarily captures the global unfolding coordinate.

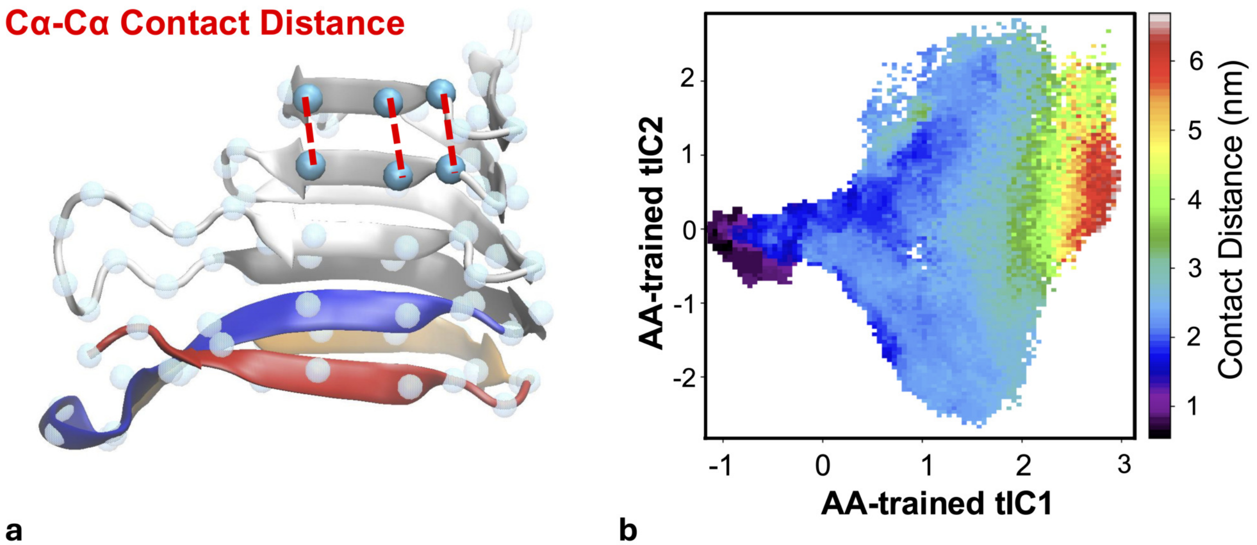


**Figure S1.** Native C$\alpha$–C$\alpha$ contacts and their relation to the unfolding coordinate. **a**, Schematic illustration of inter-rung native C$\alpha$–C$\alpha$ contact pairs in the folded $\beta$-helix. **b**, Mean C$\alpha$–C$\alpha$ distance over all native contact pairs projected onto the all-atom tIC1–tIC2 landscape, showing that tIC1 primarily tracks the global unfolding degree.

#### 2 Native Cα–Cα contact pairs used for anisotropy calculation

**Table S1.** Native Cα–Cα contact pairs used for anisotropy calculation. Listed are 47 native Cα–Cα contact pairs defined from inter-rung hydrogen-bond ladders in the folded $\beta$-helix structure. Residue numbering follows the original PDB structure of the pertactin passenger domain.

| Native Cα–Cα pair (i–j) | Native Cα–Cα pair (i–j) | Native Cα–Cα pair (i–j) |
|---|---|---|
| 379–397 | 396–414 | 424–449 |
| 380–398 | 397–415 | 425–450 |
| 381–399 | 398–416 | 426–451 |
| 382–400 | 399–417 | 427–452 |
| 383–401 | 400–418 | 438–463 |
| 384–402 | 401–419 | 439–464 |
| 388–405 | 402–420 | 440–465 |
| 389–406 | 405–424 | 441–466 |
| 390–407 | 406–425 | 442–467 |
| 391–408 | 407–426 | 442–468 |
| | 408–427 | 443–469 |
| | 412–439 | 444–470 |
| | 413–440 | 449–475 |
| | 414–441 | 450–476 |
| | 415–442 | 451–477 |
| | 416–443 | 452–478 |
| | 417–444 | 453–479 |
| | 418–445 | 454–480 |
| | 419–446 | |

#### 3 Validation of lag-time selection for coarse-grained tICA

To validate the choice of lag time for tICA-based CV construction from coarse-grained simulations, we performed a lag-time scan by projecting the CG ensemble onto the first two tICs at lag times of 0.4, 2, 4, 40, 80, and 200 ns.

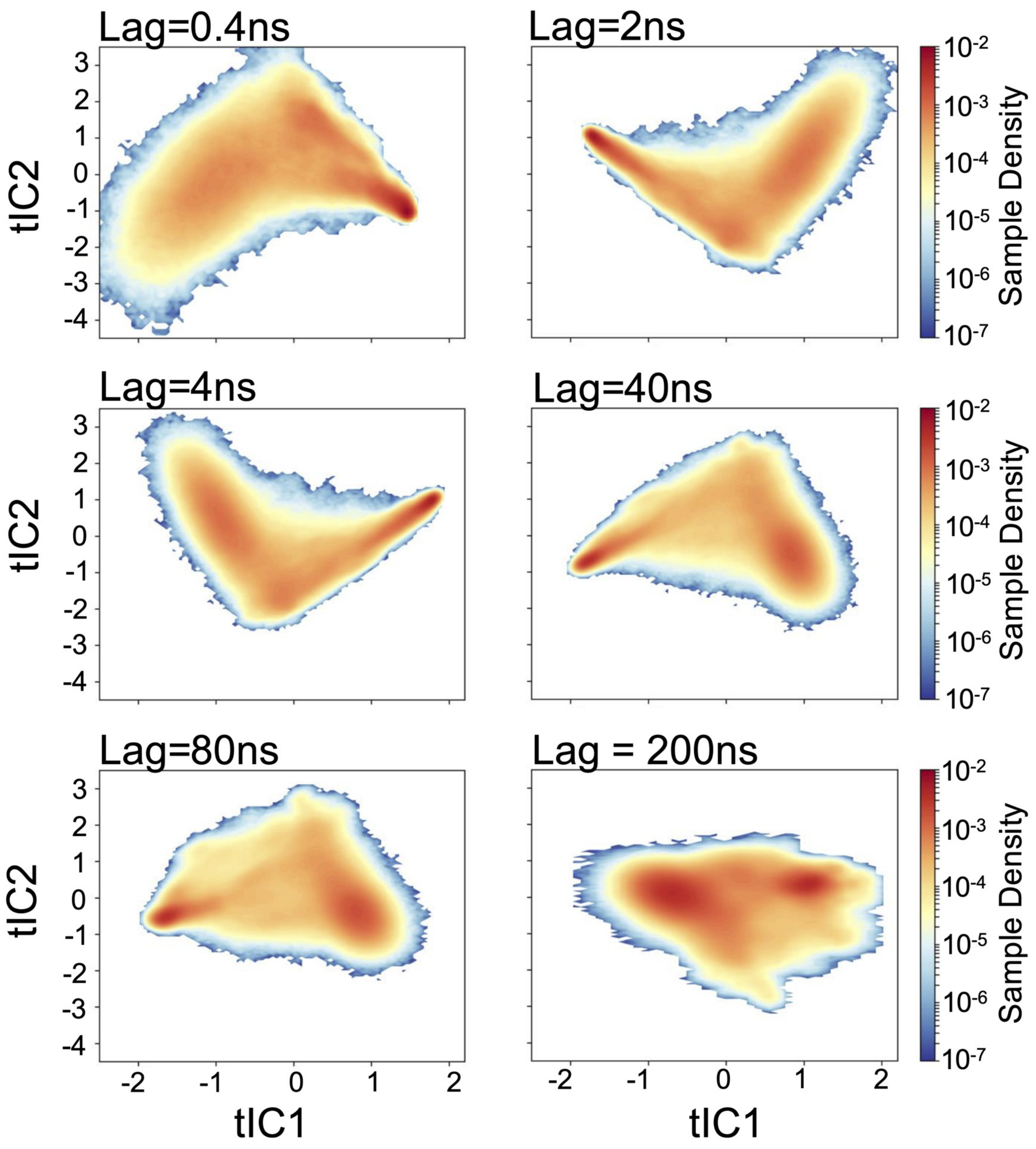


**Figure S2.** Projections of coarse-grained trajectories onto the first two tICs at six different lag times. Colors indicate sample density on a logarithmic scale.

#### 4 REUS PMF Convergence Analysis

Root-mean-square deviation (RMSD) between free-energy surfaces computed from cumulative simulation data (0 to t ns per window) and the final PMF (full 2.2 ns per window) as a function of simulation time. PMFs were anchored by setting their respective minima to zero before RMSD calculation to compare relative shapes. The RMSD drops rapidly during the first nanosecond and falls below the convergence threshold of 0.5 kcal/mol (red dashed line) after approximately 1.5 ns per window, indicating good convergence of the free-energy landscape. The final RMSD at 2.2 ns is 0.05 kcal/mol, demonstrating that the PMF has stabilized.

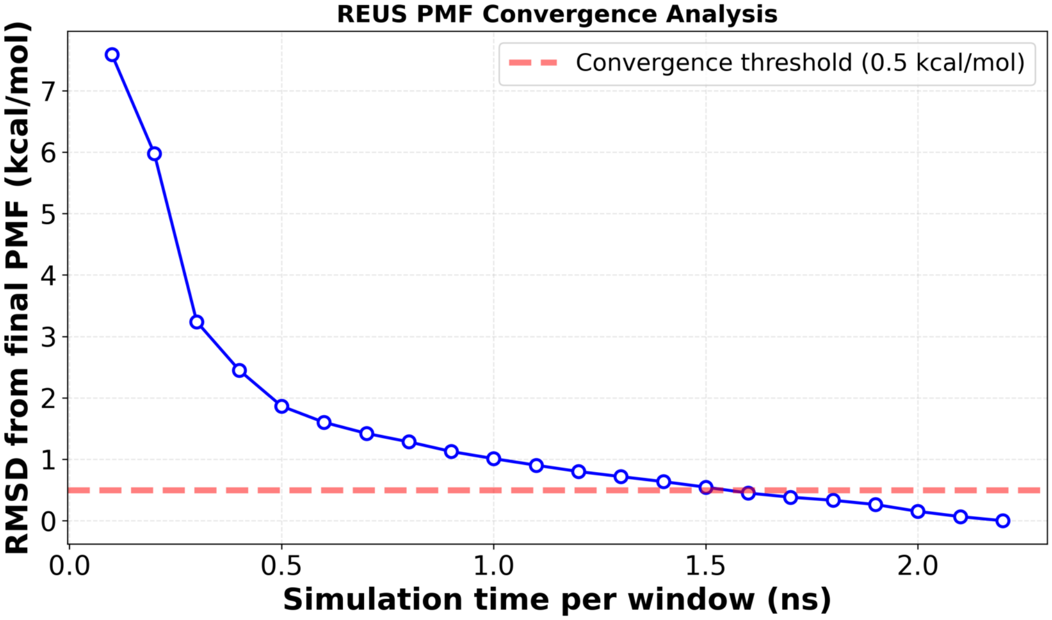


**Figure S3.** REUS PMF convergence analysis. Root-mean-square deviation (RMSD) between cumulative PMFs and the final PMF as a function of simulation time per umbrella window.
